# Hexokinase: A central player in the synergism of high-intensity intermittent exercise and every-other-day intermittent fasting regimen on energy metabolism adaptations

**DOI:** 10.1101/389668

**Authors:** Antonio Real-Hohn, Clarice Navegantes, Katia Ramos, Dionisio Ramos-Filho, Fábio Cahuê, Antonio Galina, Verônica P. Salerno

## Abstract

Visceral lipid accumulation, organ hypertrophy and a reduction in skeletal muscle strength are all signs associated with the severity of obesity related disease. Intermittent fasting (IF) and high-intensity intermittent exercise (HIIE) are natural strategies that, individually, can prevent and ameliorate obesity along with metabolic syndrome and its associated diseases. However, the combinatorial effect of IF and HIIF on energetic metabolism is currently not well understood. We hypothesized that their combination could have a potential for more than strictly additive benefits. Here, we show that two months of every-other-day intermittent fasting regimen combined with a high-intensity intermittent exercise protocol (IF/HIIE) produce a synergetic effect, preventing fat accumulation, enhancing physical performance and optimizing energy production. The IF/HIIE group presented increased glucose uptake, lower levels of serum insulin and a global activation of hexokinases in skeletal muscle, heart and liver comparing to control, IF and HIIE groups. IF/HIIE synergism led to activation of the FoF1 ATP synthase and promoted a more oxidative profile of mitochondria in observed skeletal muscle. Additionally, high-resolution respirometry of muscle fibers showed that animals in the IF/HIIE group presented characteristics suggestive of augmented mitochondrial mass and efficiency. Finally, an important reduction in serum oxidative stress markers were observed in IF/HIIE group. These findings provide new insights for the implementation of non-pharmaceutical strategies to prevent/treat metabolic syndrome and associated diseases.

## Introduction

Obesity and metabolic syndrome are both important risk factors for life threatening diseases that can target cardiovascular and hepatic systems (1, 2). The prevalence of obesity and metabolic syndrome is a reality for developed countries, which began more than two decades ago (3). Today, the incidence of obesity and metabolic syndrome is rapidly increasing in developing countries as well (4), leading to increased morbidity and mortality due to type 2 diabetes mellitus, non-alcoholic fatty liver disease, and cardiovascular disease (5, 6). Recently, global climate change was implicated in the onset of obesity and type 2 diabetes due to the negative impact of higher temperatures on energy metabolism (7, 8). This means that the overall prevalence of obesity and metabolic syndrome tends to aggravate in the next few years allied with the increased reduction in physical activity (9).

Intermittent fasting (IF) regimens and high-intensity intermittent exercise (HIIE) are two natural strategies to prevent and mitigate obesity related diseases (10, 11). An every-other-day IF regimen was recently demonstrated by Li, Xie (12) to dramatically reduce obesity, insulin resistance, and hepatic steatosis in rodents by altering the gut microbiota. Conversely, the adaptations promoted by HIIE in rodents has been demonstrated to be a direct, stimulus-driven mechanism with a global effect through the mobilization of several organs like skeletal muscles (13), liver (14) and heart (15). Both IF and HIIE approaches are appropriate treatments for obesity-related problems in humans (16, 17). Noteworthy, IF and HIIE strategies respectively resemble the evolution patterns of human diet (18), with an erratic food availability, and the high-intensity intermittent exercise analogous to hunting/gathering activities (19).

Recently, a study combining caloric restriction with HIIE revealed increased glucose uptake along with higher Glut4 and Fasn mRNA levels in skeletal muscle (20). Ever since IF can produce different adaptations compared to caloric restriction (21), we reasoned that IF associated with HIIE could have a strong synergic effect in energy metabolism through hexokinase modulation and mitochondrial reprogramming. We centered our assessment on hexokinase, as this enzyme is known to be essential to overcome the rate-limiting step of the glucose metabolism (22) and the rate-limiting step of the oxidative phosphorylation (23).

## Material and methods

### Animals and intermittent fasting protocol

All animal procedures performed received prior approval from the Animal Use Ethical Committee in the Health Science Center of the Federal University of Rio de Janeiro (Rio de Janeiro, RJ, Brazil; Protocol CEUA/EEFD06). At the beginning of the adaptation phase (one month), twenty-four, 60-day-old male Wistar rats were housed in a climate-controlled environment (22.8 ± 2.0 °C, 45–50% humidity) with a 12/12– light/dark cycle with access to food and water ad libitum. Three weeks before the beginning of the study, animals were acclimated to the experimental protocols: Two weeks under the IF regimen followed by one week with the IF regimen plus HIIE (no overload). The chow given to the animals was a standard laboratory chow Nuvilab CR-1 (Nuvital Nutrientes, Paraná, Brazil) with 22% protein, 8% fibers, and 4% fat. Animals in control (C) and HIIE groups had access to food ad libitum during all the study while those in IF and IF/HIIE groups were subjected to an every-other-day IF regimen. IF and IF/HIIE groups were provided access to food ad libitum for 24 hours that was alternated with 24 hours without food. Animals were weighed weekly in the morning before the withdrawal or reintroduction of food. Food consumption was evaluated daily and the intermittent fasting resulted in a 15% reduction in total offered calories. Importantly, at the beginning of the study, animals already had reached 90-day-old (young adults), avoiding influences of sexual maturation (30-40 days) (24) and musculoskeletal development (25) in our analyzes.

### High intensity intermittent exercise protocol and physical test

Groups HIIE and IF/HIIE performed 8 weeks of an interval swimming exercise consisting of 14 repeated 20-second swimming bouts with weight (equivalent to percent body weight) attached with 10 seconds rest between the repeats as described previously (26). Before the beginning of the study, animals were adapted one week to the aquatic conditions by performing the exercise without an overload. An initial overload of 6% of the body weight (bw) was attached to the animal during the swimming period. The load was increased by 2% bw every two weeks. The HIIE protocol was performed exclusively at Mondays, Wednesdays and Fridays during the protocol adaptation and continued during the 8 weeks of the study. This routine was employed to avoid overtraining the animals. Physical tests were used to determine changes in cardiorespiratory endurance in the animals. Each test consists of the time swimming until fatigue under a 12% bw overload, that was applied on three separate occasions: (1st) one day before day 0, (2nd) at day 28, and (3rd) at day 56 (Fig 1 and 5A).

**Fig 1.**
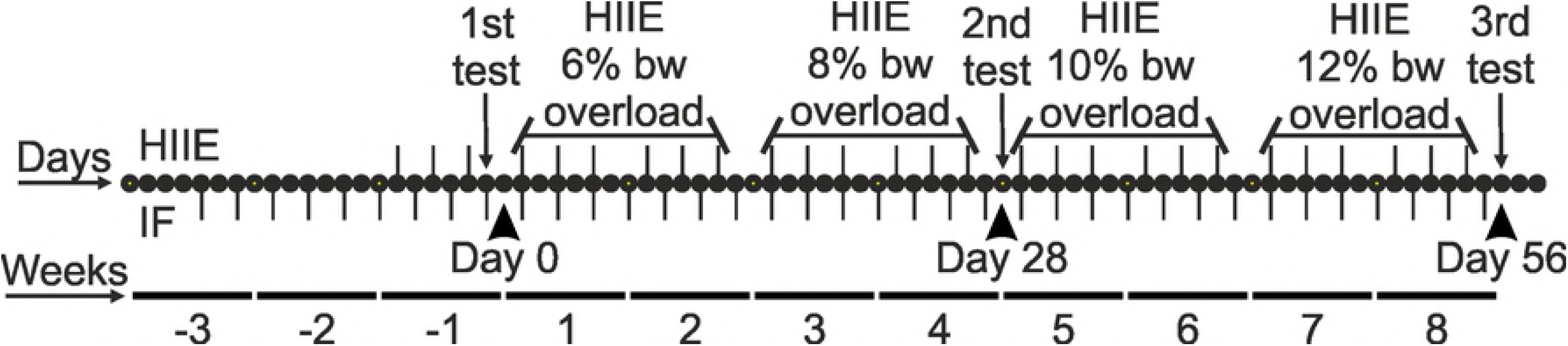
Graphical representation of the experimental design. Days (dots) and weeks (horizontal lines) of the study period (56 days in total) showing the exactly days of fasting (lower dash) and HIIE (upper dash) interventions. The load utilized for each HIIE is described above the respective days. The 1st, 2nd, and 3rd physical tests for endurance are indicated. The adaptation phase is represented by weeks with negative numbers.

### Intraperitoneal glucose tolerance tests

Animals were fasted for 12 hrs prior to the administration of an intraperitoneal injection of glucose (2 g/kg body weight). Blood samples were drawn from tail vein immediately before the glucose challenge, as well as 15, 30, 60, 90, and 120 min thereafter. Blood glucose levels were determined using an Accu-Chek glucose analyzer (F. Hoffmann-La Roche Ltd, Basel, Switzerland).

### Fasting serum insulin evaluation

Blood samples were collected in fasted animals with heparinized tubes and the serum were separated by centrifugation and kept in −80 °C. The total insulin level of the frozen serum samples was measured using an Elisa-based method by VetLab Veterinary Clinical Pathology Laboratory (Petropolis, Rio de Janeiro, Brazil).

### Tissue collection and preparation

Heart, liver, gastrocnemius muscle and visceral fat were rapidly removed from animals euthanized by decapitation 2 days after the last day of the experimental period (56 days) and weighed. For enzymatic analysis and NADH measurements, tissue was homogenized with a Potter-Elvehjem in homogenization buffer (30 mM KCl, 4 mM EDTA, 250 mM sucrose, and 100 mM Tris-HCl (pH 7.5) with the protease inhibitors aprotinin and PMSF). Homogenates were centrifuged (5000 x g for 10 min at 4 °C) to obtain the supernatants that were maintained at 4° C. For the respiration analyses, gastrocnemius muscle was minced and transferred to ice-cold BIOPS buffer (10 mM Ca^2^+/EGTA, 0.1 mM free Ca^2^+, 20 mM imidazole (pH 7.1), 50 mM K+-MES, 0.5 mM DTT, 6.56 mM MgCl_2_, 5.77 mM ATP, 15 mM phosphocreatine). Next, saponin (50 μg/ml) was added and incubated for 30 min at 4 °C, followed by a buffer exchange into ice-cold BIOPS without saponin. Samples were further incubated 2h at 4 °C prior to high-resolution respirometry experiments.

### Determination of fiber cross-sectional area

Frozen gastrocnemius histological sections (5μm) were obtained in Leica CM 1850 Cryostat (Leica Biosystems, Nussloch, Germany), fixed with 4% formal calcium and stained with the hematoxylin and eosin (HE) method. We captured images of the stained sections with Leica DM 2500 optical microscope (20 x lens) (Leica Biosystems, Nussloch, Germany). Fifty gastrocnemius myofibers from each animal of different groups were randomly selected and fiber cross-sectional area was measured using Image J 1.51n software (NIH, Bethesda, MD, USA). A total of 150 myofibers were plotted for each group to provide a reasonably reliable estimate of the total fiber number (27).

### Hexokinase activity

The method to measure enzymatic activity was performed at 37 °C in buffer containing 4 mM MgCl_2_, 50 mM Tris-HCl (pH 7.5), 20 mM glucose (0.1 mM glucose for liver), 4 mM ATP, 1 U/ml G-6PDH, 0.5 mM β-NADP+, and 0.1% Triton X-100 with a protein concentration of 0.05 mg/ml. The absorbance at 340nm was acquired every 30 seconds for 30 minutes and the enzymatic activity was calculated using a molar extinction coefficient of 0.00622 uM^-1^ cm^-1^ for NADPH.

### Oxidative profile measurements

The oxidative profile of skeletal muscle was determined through the measurement of its mitochondrial NADH content through a fluorescence assay. Briefly, 140 μg of protein from a muscle sample homogenate derived from each animal was placed into a 98 well plate and excited at 340 nm. The emission at 450nm was measured in a Spectramax Paradigm (Molecular Device, Sunnyvale, CA, USA). The assay was repeated three times and the fluorescence values were plotted as arbitrary numbers.

### High resolution respirometry

Respiration measurements were performed on fiber bundles in 2 ml of mitochondrial respiration medium 05 (110 mM sucrose, 60 mM potassium lactobionate, 0.5 mM EGTA, 3 mM MgCl_2_, 20 mM taurine, 10 mM KH_2_PO_4_, 20 mM HEPES (pH 7.1), 2 mg/ml BSA). O_2_ consumption was measured using the high-resolution Oxygraph-2k system (Oroboros Instruments GmbH, Innsbruck, Austria). The results were normalized to the wet weight of the permeabilized fiber bundles. All the experiments were performed at 37 °C in a 2 ml chamber. Mitochondria membrane permeability was tested by the addition of 10 μM cytochrome c. No greater than a 10% increase in oxygen consumption was observed. Multi-substrate titrations, respiratory states and respiratory control ratio calculations were performed. State 3 was measured after addition of complex I substrate, complex II substrate, and ADP. State 4o was measured subsequently to State 3 after addition of oligomycin to mimic State 4. State uncoupled was measured in the sequence through addition of FCCP. RCR was calculated by State 3 divided by State 4o (28).

### FoF1 ATP synthase activity assay in skeletal muscle

ATP synthase activity was extrapolated from ATP hydrolysis (ATPase) activity (29). Briefly, 50 μg of protein from gastrocnemius homogenate from each animal was used to measure ATPase activity in 1 mM ATP, 5 mM MgCl_2_, and 50 mM Tris (pH 8,5) buffer with the presence or absence of 5 mM sodium azide at 37 °C. After TCA precipitation (20% w/v) and centrifugation (3000 x g for 15 minutes at 4 °C) the resultant supernatant was collected and combined with ammonium molybdate and Fiske-Subbarow reducer. The absorbance was measured at 660 nm.

### Protein oxidation

Protein oxidation levels were measured using the protein carbonyl content method (PCC), as previously described (30). Briefly, the blank sample was mixed with 2.5 N HCl and the other with 2,4-dinitrophenylhydrazine (freshly prepared in 2.5 N HCl) and the resulting solutions were incubated in dark for 1 h at RT with intermittent vortexing (every 15 min), with subsequent addition of 10% TCA (w/v). After centrifugation, the pellet was washed once with 10% TCA and three times with ethanol: ethyl acetate (1:1 v/v). The resulting pellets were suspended in 5 M urea (pH 2.3), incubated at 37 °C for 15 minutes and centrifuged at 15000 x g for 5 minutes. The resulting supernatant absorbance was determined at 370 nm, and results were expressed as nmol carbonyl / mg protein.

### Lipid peroxidation

Lipid peroxidation levels were measured using thiobarbituric acid method (TBARS), with minor modifications of the technique previously described (31). Serum samples were diluted in 100 mM sodium phosphate buffer (pH 7.4), 1:3 (v/v), with subsequent addition of cold 10% TCA and kept on ice for 15 minutes. After, samples were centrifuged at 2200 x g for 15 min (4 °C) and to the resultant supernatants were added equal volumes of 0.67% thiobarbituric acid (w/v) followed by water bath (95 °C) incubation for 2 h. After cooling, the absorbances were read at 532 nm in 96-well plate reader, Spectra Max Paradigm (Molecular Devices, California, United States). Results were expressed in μM malondialdehyde (MDA).

### Statistical analysis

Comparisons were performed using two-way ANOVA with multiple comparison test. Data are presented as mean ± standard deviation (SD) and P values < 0.05 were considered significant. All statistical analyzes were performed using Prism 7.0 (Graph Software Inc., La Jolla, CA, USA).

## Results

### IF/HIIE protocol prevented weight gain and increased muscle crosssectional area

To determine the adaptive changes on energetic metabolism and physical performance induced by IF, HIIE, and their combination, the three regimens were imposed on age-matched young adult Wister rats over 8 weeks (Fig 1). Over the course of the experimental conditions, the weight of the animals was tracked weekly and plotted in a curve (Fig 2A). The cumulative increase in the weight of animals on an ad libitum diet (C and HIIE groups) is very apparent and the prevention of excessive weight gain in animals under IF protocol (IF and IF/HIIE groups) is evident; other groups observed a similar effect (12, 18). To explore the possible origin of the observed differences in the weight of the groups, organs and tissues were collected and weighed at the end of the study. Initially, we weighed the brown adipose tissue (Fig 2B) and the visceral fat (Fig 2C) of the animals. Our results were similar to those observed in Li et al., with no variation in brown adipose tissue mass and an evident reduction in visceral fat mass in groups under IF protocol (IF and IF/HIIE) (12). We also observed a reduced total mass of the liver (Fig 2D) and heart (Fig 2E) in animals under the IF protocol (IF and IF/HIIE groups). A reduction in skeletal muscle (gastrocnemius) total mass (Fig 2F) was observed in the IF group and an evaluation of the cross-sectional area (CSA) confirmed the indications of atrophy (Figs 2F and 2G). In contrast, the measurement of CSA from the IF/HIIE group was indicative of hypertrophy of the myofibers followed to a lesser extent by the HIIE group (Figs 2F and 2G).

**Fig 2.**
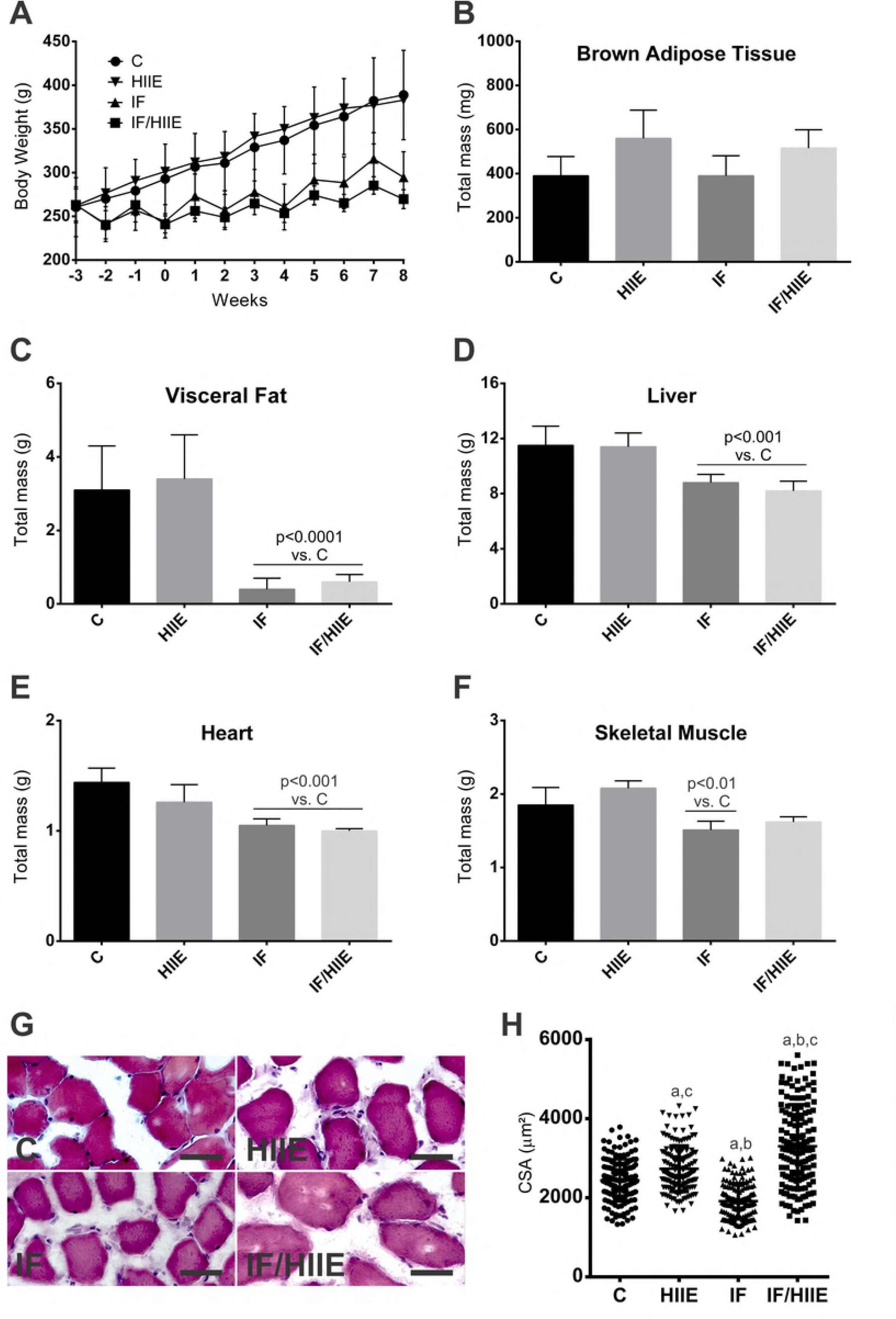
IF/HIIE prevented weight gain, adiposity and increases skeletal muscle cross-sectional area. (A) The body weight curve from the weekly weight weekly obtained in the morning before the withdrawal or reintroduction of food. Tissues and organs were collected from animals 2 days after day 56, end of the study period, and immediately weighed. (B) Brown adipose tissue weight; (C) visceral fat weight; (D) liver weight (intact organ); (E) heart weight (intact organ); (F) skeletal muscle weight (gastrocnemius); (G) cross-sectional area histological profile (myofibers were stained with HE; 50 μm bar) and (H) quantification (dot blot representation with average and standard deviation bars of 150 myofibers per group). p < 0.05; a vs. control, b vs. HIIE, and c vs. IF.

### Serum glucose uptake is integrated with hexokinase activity in IF/HIIE group

To investigate how IF and HIIE could impact the systemic glucose availability and metabolism, we measured the fasting blood glucose levels, rate of glucose uptake and fasting serum insulin levels (Fig 3). All groups presented similar values for fasting blood glucose (Fig 3A). Initially, the glucose tolerance test also showed similarities between groups (Fig 3B). However, an analysis of the area under the curve (AUC) revealed that the IF/HIIE group presented a significantly faster glucose uptake (Fig 3C). Glucose uptake allied to fasting serum insulin levels that could present a predictive factor of insulin sensitivity (32, 33). For this reason, we measured the fasting insulin level of the different groups (Fig 3D) and we observed an important reduction for the IF/HIIE group followed by the IF group.

**Fig 3.**
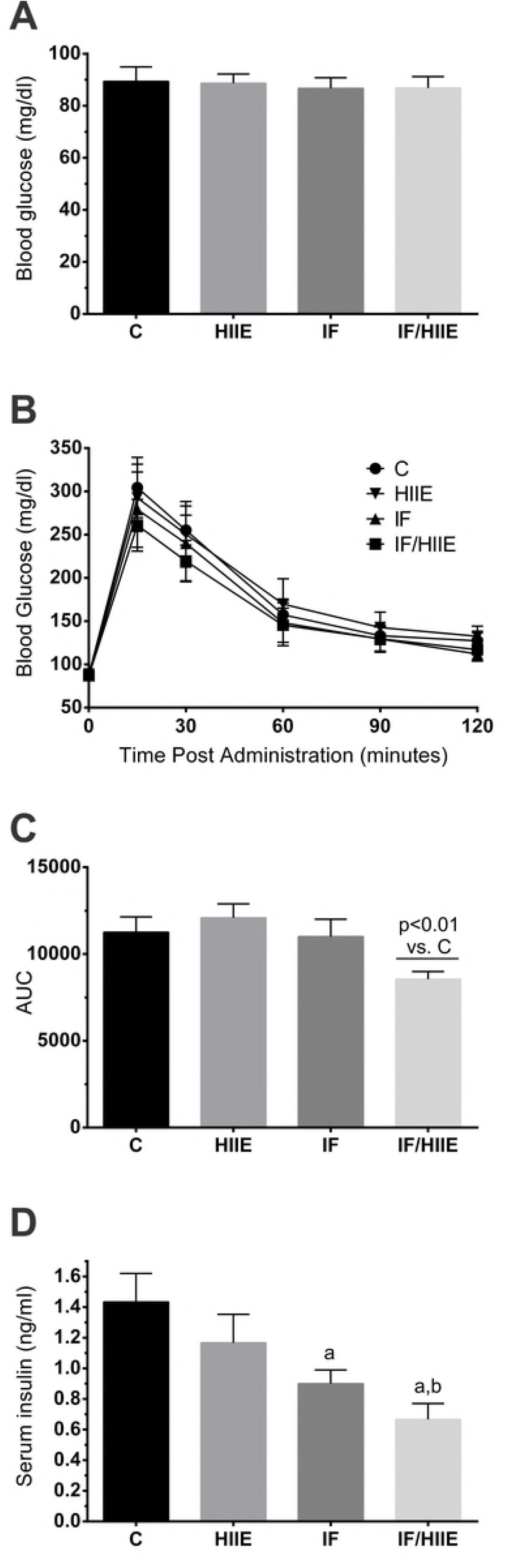
IF/HIIE activates glucose uptake and reduces fasting serum insulin levels. (A) Serum blood glucose levels. (B) Glucose tolerance test. Animals were fasted for 12 h prior to the administration of an intraperitoneal injection of glucose (2 g/kg body weight). (C) AUC calculated from glucose tolerance test curve (B). (D) Insulin serum levels of fasted animals were measure using an Elisa-based method. p < 0.05; a vs. control and b vs. HIIE.

According to the literature, sugar transport has been integrated with hexokinase (HK) activity in a cellular model (34). To test if this mechanism could explain observations in a more complex system, we hypothesized that increased blood glucose uptake could be integrated to an increased HK activity in multiple organs (Fig 4). In the liver, we observed that the IF/HIIE group had the highest HK activity followed by HIIE and IF groups, which showed similar activity values (Fig 4A). In heart, both IF and IF/HIIE groups showed increased HK activity (Fig 4B). In skeletal muscle, the IF/HIIE group had the highest HK activity (6-fold increase) followed by HIIE group (3-fold increase). Taken together, the HK activity data from all analyzed organs suggest that IF/HIIE group possess the highest values for HK activity.

**Fig 4.**
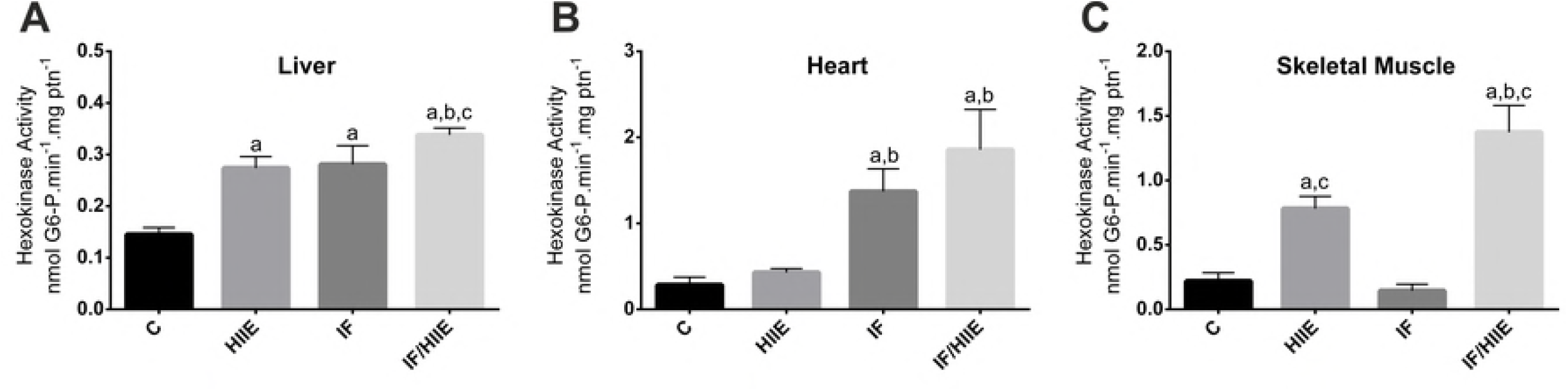
Synergic effect of IF/HIIE on HK activity. Enzymatic activity was calculated from NADPH production. (A) HK activity in liver. (B) HK activity in heart. (C) HK activity in skeletal muscle (gastrocnemius). p < 0.05; a vs. control, b vs. HIIE, and c vs. IF.

### IF/HIIE synergic effect in physical activity and energy production

The effect of HIIE promoting physiological adaptation in skeletal muscle is well described (14, 35). However, the effect of IF combined with HIIE in physical performance is not, although some reports that employed endurance training suggest a possible synergetic effect: I) Rodriguez-Bies et al. combined IF protocol with endurance exercises and observed a consistent increase in beta-oxidation, lactate production, and mitochondria content in gastrocnemius with a modest effect in physical performance in animals submitted to both protocols compared to the control group (36). II) Moraes et al. showed preserved muscle mass in animals submitted to an IF protocol allied to endurance exercises (37). However, the physical capacity of these animals was not evaluated in the latter. To investigate a possible physical improvement promoted by IF and/or HIIE along the experimental procedures, we submitted all groups to a physical test (PT) on three days (Fig 5A): 1st) one day before day 0; 2nd) at day 28; 3rd) at day 56 (end of the IF and HIIE protocols). The 3rd test revealed a higher performance of animals in the HIIE and IF/HIIE groups that was approximately 90% and 180% in comparison to control, respectively. The possibility that the weight of the animals from IF and IF/HIIE groups contributing to the outcome of the PT was eliminated since no correlation was observed between swimming time and weight of the animals in any of the PTs (S1 Fig).

**Fig 5.**
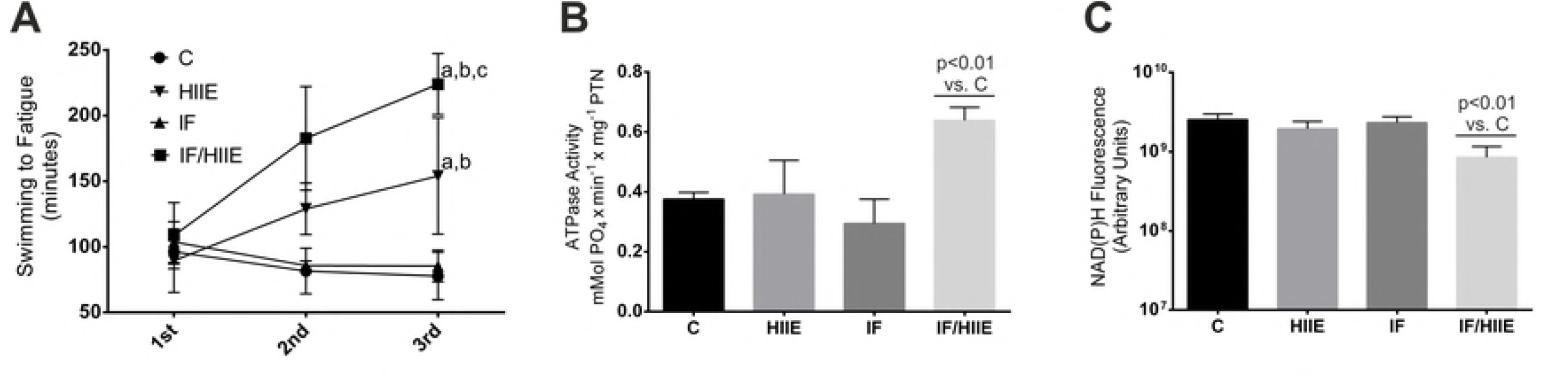
Physical test and energy production. (A) Physical test. The physical endurance was measured using swim until fatigue test (under a 12% bw overload). The tests were applied on three separate occasions: (1st) one day before day 0, (2nd) at day 28, and (3rd) at day 56. (B) FoF1 ATP synthase activity was estimated through the ATPase activity of the enzyme. (C) NAD(P)H autofluorescence was measured directly in fresh muscle homogenates (gastrocnemius). The excitation and emission wavelengths were 340 nm and 450 nm, respectively. p < 0.05; a vs. control, b vs. HIIE, and c vs. IF.

Furthermore, to explore any possible adaptation that granted IF/HIIE group the best results in the physical test, we measured the activity of the FoF1 ATP synthase in the skeletal muscle of these animals (Fig 5B). We observed an approximately 50% increase in FoF1 activity in IF/HIIE group compared to the other groups. According to the literature, both IF (36) and HIIE (38, 39) alone could promote adaptations in mitochondria. To access the mitochondria oxidation profile, we measured the NAD(P)H content in skeletal muscle of the groups (Fig 5C). Within mitochondria, NADH maintains a supply of protons for the redox couples of the electron transport chain. Blockade of the electron transport slows the rate of NADH oxidation and raises NADH/NAD ratio; lower NADH/NAD ratio should be accompanied by higher NADH oxidation and improved electron transport (40). As it is not possible to distinguish between the fluorescence of NADH and NAD(P)H or between cytosolic and mitochondrial nucleotides, the fluorescence signal was referred to as NAD(P)H. The bulk of the measurement was assumed to be from mitochondria since cytosolic NADH and NAD(P)H contribute in general less than 20% of the signal under these conditions (41). Only the IF/HIIE group presented a more oxidative profile, ratifying ATP synthase enhancement observed in IF/HIIE group.

### Synergic effect of IF/HIIE in muscle fiber mitochondria respiratory states

To investigate how the skeletal muscle mitochondrial respiratory complexes could be affected by IF and/or HIIE protocols, we measured the different respiratory states and the mitochondria respiratory control rate (RCR) (Fig 6). RCR is an reliable indicator of mitochondria function, as high RCR usually indicates healthy mitochondria and low RCR usually indicates mitochondrial dysfunction (42). Initially, we analyzed muscle fiber mitochondria oxygen consumption in the presence of complex I substrate, complex II substrate, and ADP (State 3). We observed a higher O_2_ flux rate related to ATP production in IF/HIIE group, followed by both HIIE and IF groups individually, in comparison to the C group (Fig 6A). In the sequence, we added ATP synthase inhibitor (State 4o). HIIE group presented a higher O_2_ flux (Fig 6B), indicative of increased proton leakage and/or extra-mitochondrial O2 consumption. In contrast, IF and IF/HIIE groups presented a lower O2 flux in comparison to C group (Fig 6B). Next, FCCP was added to uncouple O_2_ flux to ATP production (State uncoupled). The IF/HIIE group presented higher values in comparison with the other groups (Fig 6C), indicative of an augmented number of respiratory complexes and/or mitochondrial mass. The RCR calculated values were greatest in the IF/HIIE group followed by the IF group, both of which were significantly greater than the values of C and HIIE groups that were nearly equal (Fig 6D).

**Fig 6.**
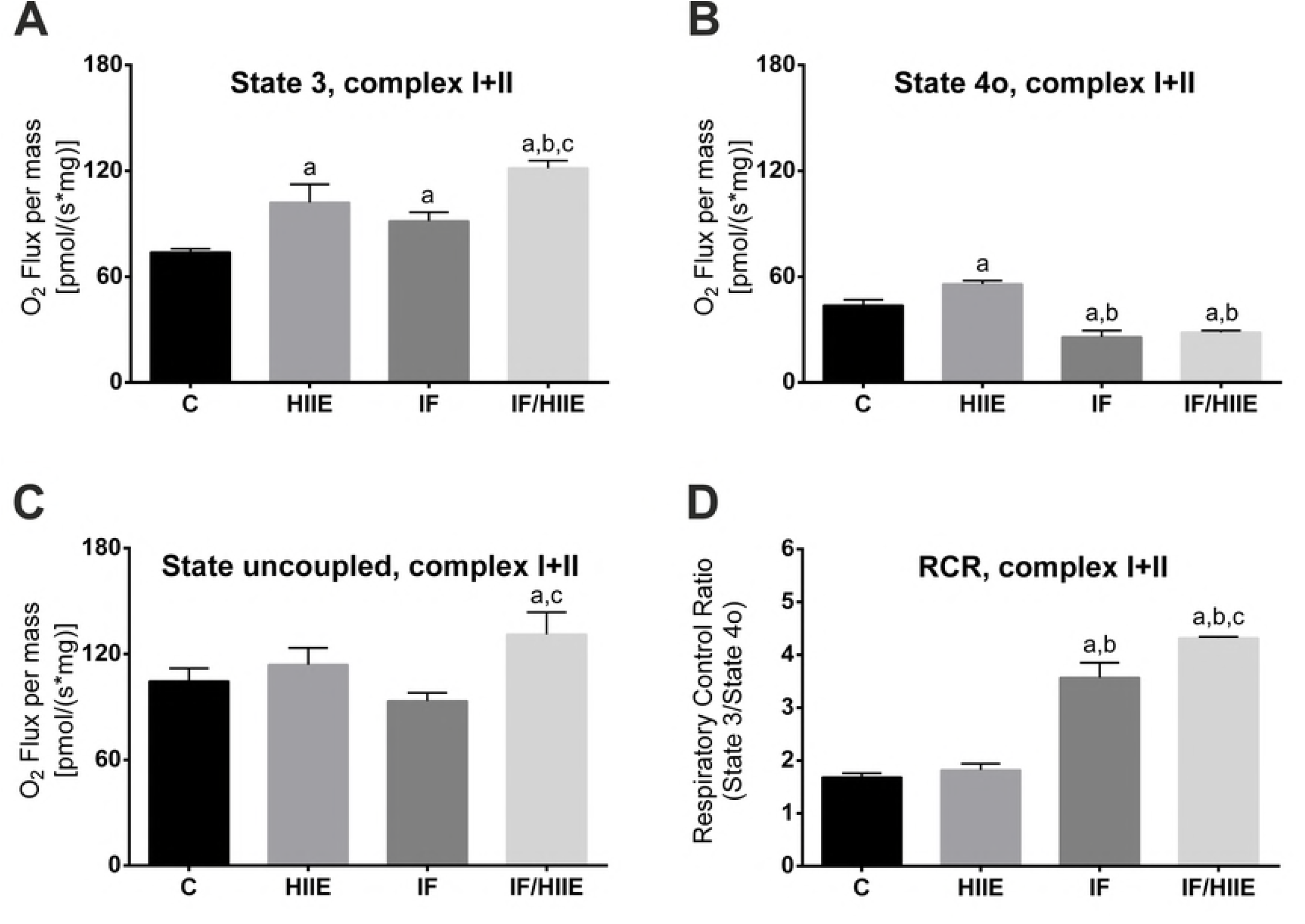
IF/HIIE promotes mitochondria activation in permeabilized skeletal muscle fiber bundles. Respiration measurements were performed on gastrocnemius fiber bundles using a high-resolution respirometry. The results were normalized to the wet weight of the permeabilized fiber bundles. (A) State 3 O_2_ flux. State 3 was measured after addition of complex I and II substrates and ADP. (B) State 4o O_2_ flux. State 4o was measured subsequently to the State 3 after addition of oligomycin to mimic real State 4 (depletion of ADP). (C) State uncoupled O_2_ flux. State uncoupled was measured in the sequence of state 3 and state 4o through addition of FCCP. (D) RCR calculation. RCR was calculated by State 3 divided by State 4o. p < 0.05; a vs. control, b vs. HIIE, and c vs. IF.

The results above are indicative of an improved O2 flux and ATP production coupling in the IF/HIIE and IF groups. The respiratory profile of the IF/HIIE group resembles data from a different group that observed an augmented mass of mitochondrial in skeletal muscle using a dissimilar exercise protocol (36). Finally, the greater RCR value agrees with a more oxidative profile and active FoF1 synthase that was observed exclusively in IF/HIIE group.

### Overall oxidative stress markers are reduced in IF/HIIE group

High intensity exercise was clearly demonstrated to negatively modulate the muscular redox state with a resultant increase in lipid peroxidation (43). However, the adoption of HIIE models was also shown to induce positive effects in muscular physiology (reviewed in MacInnis and Gibala (44)). We hypothesized that in addition to local effects promoted by HIIE in skeletal muscle, it could also affect the overall redox state with a modulation in oxidative stress markers and, in combination with IF, could possibly generate a synergistic effect.

To investigate the effect of IF and HIIE in the overall redox state, we measured the serum level of the oxidative stress damage markers lipid peroxidation and protein oxidation through malondialdehyde (MDA) quantification and protein carbonyl content, respectively. We observed a strong reduction in MDA levels only in IF/HIIE group (Fig 7A) showing a synergic effect in prevention of lipid peroxidation. Furthermore, IF/HIIE group presented lower levels of serum protein oxidation followed by HIIE group (Fig 7B). The IF group presented similar values as the control group for the oxidative markers (Figs 7A and 7B) indicative of an absence of an increase in the oxidative stress protection in serum.

**Fig 7.**
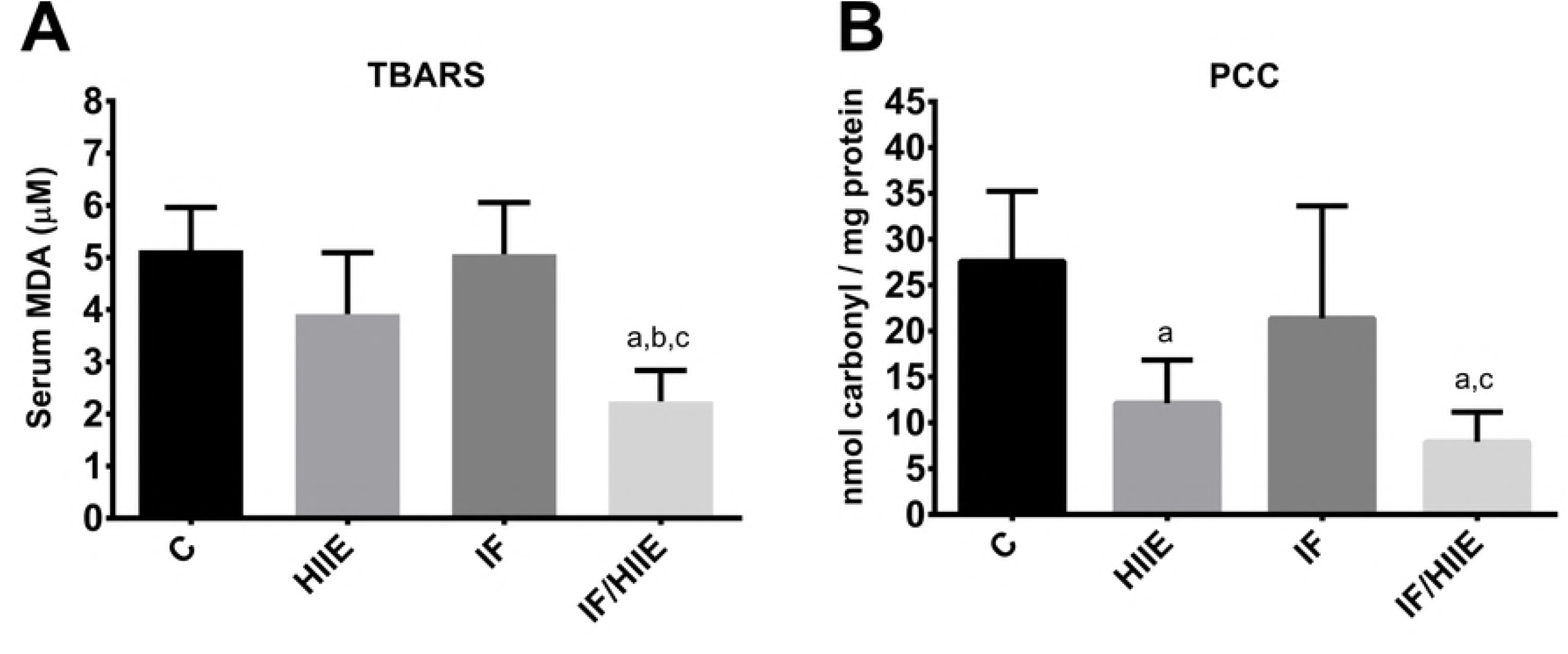
IF/HIIE group presented lower levels of serum lipid peroxidation and protein oxidation. Oxidative damage markers were measured in the serum through quantification of (A) lipid peroxidation (MDA) and (B) protein oxidation (PCC). p < 0. 05; a vs. control, b vs. HIIE, and c vs. IF.

## Discussion

Over the last few years, IF (10, 45) or HIIE (46, 47) protocols were individually evaluated aiming the potential to promote energy metabolism and physiologic adaptations. Different combinations of fasting regimens with a variety of exercise protocols were also employed (48), however, none of these studies has examined the interaction between IF and HIIE. Only recently, researchers have begun to recall the mechanism underneath IF and HIIE adaptations. One of the mechanisms that has been proposed is that IF shapes the microbiota in the gut and the adaptions to the microbiota can induce the same modifications when transplanted to a germ-free animal (12). For HIIE, a proteomic change has been proposed for skeletal muscle cells that are especially noticeable in mitochondria and ribosome protein profiles (49). These discoveries shed new light on IF and HIIE that motivated us to investigate a potential synergic effect from the combination of these strategies. To test this hypothesis, young adult Wistar rats (3-month-old) were submitted to 2 months of IF and/or HIIE (Fig 1) with an additional month of adaptation prior to the start of the study.

Initially, we observed that the IF fasting protocol promoted a pronounced protection against the weight gain observed in groups with food offered ad libitum (Fig 2A). According to the literature, IF promotes activation of brown adipose tissue (12), however, no noticeable difference in size or weight of brown adipose tissue was observed in any of the groups (Fig 2B). In contrast, IF and IF/HIIE groups presented an expressively low visceral fat mass (Fig 2C). We also weighed the liver and heart of the animals, as these organs can suffer lipid accumulation in some diseases (50). Animals in both the IF and IF/HIIE groups had smaller livers (Fig 2D) and hearts (Fig 2E) suggesting that the normal diet offered ad libitum can induce aspects of obesity in young adult rats. To investigate the possibility of muscular atrophy promoted by lack of physical activity combined with reduced food offer in IF group, the weight (Fig 2F) and the CSA of gastrocnemius were evaluated (Figs 2G and 2H). We observed lower weight and CSA of gastrocnemius from IF group comparing to C group, however, the IF group presented similar results to the C group in the physical tests (Fig 5A). Surprisingly, the IF/HIIE group showed an increase in myofibers CSA (Fig 2H), statistically higher than CSA measured in HIIE group, indicating a morphological adaptation promoted by the synergy of our IF/HIIE protocol. The latter may be explained due to a possible negative regulation on activin/myostatin signaling promoted by the IF protocol that could potentiates HIIE muscle adaptation. This hypothesis was raised based on recent data demonstrating that the inhibition of activin/myostatin signaling in mice results in skeletal muscle hypertrophy (51). Furthermore, in humans, acute fasting is known to promotes reduction of activins circulating levels (52–54), and possibly, the successive cycles of fasting promoted by the IF protocol could exacerbate this effect. However, the effect of IF on activin/myostatin signaling was not the aim of our study and should be proper investigated.

To evaluate the effects of IF and HIIE protocols on glucose metabolism, core of energetic processes (55), we started measuring the fasting blood glucose (Fig 3A), which showed similar values for all groups. Next, we measured glucose uptake via glucose tolerance test (Fig 3B) with AUC analysis (Fig 3C) and we detected a significantly faster uptake of glucose in IF/HIIE group, similarly to observed by other groups (20, 37). Additionally, we measured the fasting serum insulin levels and we observed a significant decrease of insulin levels in IF/HIIE group (53% reduction), followed by a smaller reduction in insulin levels in IF group (37 % reduction). Lower levels of circulating insulin were also observed in mice over-expressing follistatin-like 3 (56), a natural blocker of activin/myostatin signaling, and possibly these groups (IF and IF/HIIE) may have higher levels of follistatin-like 3, that could corroborate the effect proposed above for IF in muscle hypertrophy through activin/myostatin signaling modulation.

To investigate the consequence of the observed increased in glucose uptake combined with low levels of circulating insulin in the downstream of glucose metabolism within the cellular milieu, we measured the activity of the first enzyme of the glycolysis, HK (Fig 4), whose activity is integrated with sugar transport (34). We observed a synergic effect from the combination of IF with HIIE protocols in liver HK with an ~130% increase versus the ~90% observed for both IF and HIIE alone (Fig 4A). In skeletal muscle HK, we observed a 3-fold greater HK activity in HIIE group compared to controls that increased to 6-fold with the combination of HIIE with IF (Fig 4C). Remarkably, the IF protocol alone showed no promotion of any adaptations in muscle HK activity, yet its combination with HIIE doubled the HK activity in the IF/HIIE group. In contrast, the activation of HK activity in the heart showed an association only with the inclusion of the IF protocol (IF and IF/HIIE groups; Fig 4B). This probably reflects an effect of high acetate serum levels as described by Li, Xie (12) in work that used a similar IF protocol. Furthermore, they also observed an increased level of circulating lactate however we discard the influence of intracellular lactate since lactate were demonstrated to reduce hexokinase activity (57).

The HK activity is closely connected with mitochondrial activity, since the muscular isoform of this enzyme can bind to the mitochondrial membrane through VDAC (58) that can positively modulate the activities of both (59). To identify a possible consequence from the strong activation of HK in muscle of the IF/HIIE group, we initially compared the results of the physical endurance test (Fig 5A). The advantage gained in performance by the combination of the IF and HIIE protocols group was patently observable. To more deeply explore the possible molecular adaptations promoted by the synergism of IF and HIIE, the mitochondria FoF1 ATP synthase in gastrocnemius muscle was measured in the groups (Fig 5B). The IF/HIIE protocol promoted an ~50% increase in FoF1 ATP synthase compared to the other groups. This effect is in agreement with the muscle HK results observed in IF/HIIE group (Fig 3C). To further confirm the possible mitochondrial activation, we measured the NAD(P)H content of the gastrocnemius of the groups (Fig 5C). Only the IF/HIIE group showed a significant reduction in the NAD(P)H content suggesting the presence of more oxidative mitochondria (60). Recently, a study combining intermittent food deprivation with an endurance running protocol for one month observed an increase in mitochondrial DNA and complex proteins along with mRNA for NRF1, NRF2 and TFAM (61). This evidence from the literature allied with the increased CSA (Figs 2G and 2H), higher FoF1 ATP synthase activity (Fig 5B), and the more oxidative mitochondria (Fig 5C) observed in IF/HIIE group that indicate a synergic effect of the protocols leading to mitochondria biogenesis. This proposed increase in mitochondrial mass could explain the increased HK activity (Fig 4). Furthermore, Zhang et al. showed recently that c-Src plays an important role in HK activity in cancer cells reducing the K_m_ the Vmax, and the tumor growth is dependent of HK. Moreover, this tyrosine kinase is known to be important during early embryonic stages as well as in ensuing differentiation processes (62, 63). Altogether, these tissue developmental effect of c-Src could explain the observed increase in muscle CSA (Figs 2G and 2H) and the increase in HK activity (Fig 4) but need to be further evaluated.

According to the literature, IF combined with diverse exercise protocols could increase AMPK and SIRT1 signaling in muscles (64) and possibly increase the mitochondrial mass (36). To directly test the effect of IF and/or HIIE on mitochondrial metabolism, we measured the O2 flux in muscle fibers (gastrocnemius) of the different groups with a high-resolution respirometry (65) (Fig 6). The results showed that the IF/HIIE group presented higher values in both the State 3 (Fig 6A) and State uncoupled (Fig 6C), which are indicative of a higher mitochondrial mass and are in agreement with previous data from another group (36). Moreover, the IF and IF/HIIE groups possessed lower values for State 4o (Fig 6B) that suggests a lower proton leak and/or extra-mitochondrial O2 consumption. The RCR calculation reinforces the idea of a more coupled mitochondria in the IF/HIIE group. The IF group also demonstrated an increased in RCR, however, even with more coupled mitochondria, this group failed to demonstrate higher levels of FoF1 ATP synthase activity (Fig 5B) probably due to a low mitochondria mass. Altogether, the activation of both HK and ATP synthase along with the more oxidative and coupled mitochondria observed in IF/HIIE group could lead to a reduction in reactive oxygen species (66) and an increased global antioxidant activity. To evaluate the possible effect of IF/HIIE in global antioxidant activity we measure the levels of serum oxidative stress markers (Fig 7). We observed that the IF/HIIE group presented with lower levels for lipid peroxidation (Fig 7A) and protein oxidation (Fig 7B), which agree with our previous hypothesis. Finally, though the data revealed here, the IF/HIIE protocol reveals a reasonable approach to be employed for translational investigation in humans, especially when considering the evolutionary-based proposal to combine IF and HIIE to mimics erratic food behavior and occasional high-intense energy demand of the gathering/hunting activities of our ancestors to improve metabolism. Different factors influence exercise tolerance in human population (specially to age and health status) and HIIE protocol must be adapted accordingly.

## Acknowledgments

The authors would like to thank Dr. David William Provance, Jr. for his review of the English. We would also like to thank Prof. Mauro Sola-Penna for all the support with the sample preparations and experiment implementation.

## Author contributions

ARH, CN and VPS designed research; CN, KR and DRF performed research; ARH, CN, KR, FC, DRF, AG and VPS analyzed data; ARH, AG and VPS contributed with new reagents or analytic tools; and ARH, FC and VPS wrote the article.

## Supporting information

**S1 Fig Analysis of the correlation between animal weight and the swimming time to fatigue in each of the three Pts.** Individual values correlating weight and swimming time to fatigue for every animal obtained in each PT day were plotted and the correlation were analyzed and indicated in the figures.

